# Limits to a classic paradigm: Most transcription factors regulate genes in multiple biological processes

**DOI:** 10.1101/479857

**Authors:** Daniela Ledezma-Tejeida, Luis Altamirano-Pacheco, Vicente Fajardo, Julio Collado-Vides

## Abstract

Transcription factors (TFs) are important drivers of cellular decision-making. When bacteria encounter a change in the environment, transcription factors alter the expression of a defined set of genes in order to adequately respond. It is commonly assumed that genes regulated by the same TF should be involved in the same biological process. Examples of this are methods that rely on coregulation to infer function of not yet annotated genes. We have previously shown that only 21% of TFs regulate functionally homogeneous genes based on the proximity of their catalyzed reactions in the metabolic network. Here, we provide more evidence to support the claim that a one TF/one process relationship is not a general property. We show that the observed functional heterogeneity of regulons is not a result of the quality of the annotation of regulatory interactions, or the absence of protein-metabolite interactions, and is also present when function is defined by Gene Ontology terms. Furthermore, the observed functional heterogeneity is different from the one expected by chance, supporting the notion that it is a biological property. To further explore the relationship between transcriptional regulation and metabolism, we analyze 5 other types of regulatory groups and identify complex regulons (i.e. genes regulated by the same combination of TFs) as the most functionally homogeneous, which is supported by coexpression data. Whether higher levels of related functions exist beyond metabolism and current functional annotations, remains an open question.

## INTRODUCTION

Transcription factors (TFs) are important drivers of bacterial decision-making. They convey environmental cues into the gene expression machinery by binding to specific metabolites. In turn, they promote the recruitment, or dismissal, of the RNA polymerase in a defined set of promoters. Classical examples such as LacI (Pardee, Jacob, and Monod 1959) or TrpR (Gunsalus and Yanofsky 1980) introduced the notion that individual TFs mediated defined responses, such as lactose utilization or tryptophan biosynthesis. The assumption followed that all the genes regulated by the same TF (termed regulon (Maas and Clark 1964; Neidhardt 1987)) were involved in the same biological process. Nowadays, there are various examples of widely studied TFs that are involved in more than one biological process, but they are considered special cases termed “global regulators” (Martínez-Antonio and Collado-Vides 2003), and the wide variety of processes in which they take place is easily rationalized by the large size of these regulons. Moreover, it is common to refer to TFs by the biological process they are involved in, such as “regulator of response to oxidative stress”.

The increasing availability of data on regulatory interactions and gene expression has allowed the development of several algorithms that predict new regulatory interactions (Wu and Li 2008; Göhler et al. 2011; Fitzgerald, Bonocora, and Wade 2014), new gene functions (Wu and Li 2008; Brohée et al. 2011; Liu et al. 2016), or describe modules of coregulated genes (Lemmens et al. 2006; Pérez-Rueda et al. 2015). Given the complexity of the data, the methods rely on several assumptions. Among them are: (1) genes regulated by the same TF will be involved in the same biological process, (2) genes regulated by the same TF are expected to be coregulated. These suppositions are mostly used to, either enrich the results with true positives, or evaluate the efficiency of the method. However, there is no systematic study to date that quantitatively analyses whether this is a general property of TFs, or an attribute of a few classical examples.

We have shown before that local TFs have a gradient of functional homogeneity, and that less than a quarter of TFs have a one-to-one correspondence with biological processes, in terms of the connectivity of the metabolic subnetworks directly affected by the regulatory action of the TF (Ledezma-Tejeida, Ishida, and Collado-Vides 2017). Here, we show that the observed gradient is not a result of missing metabolite-protein interactions, or low-confidence interactions, and is different from what would be expected by chance. We support that the gradient is a biological property by reassessing the functional homogeneity of each regulon using Biological Process Gene Ontology terms. In order to find an alternative to general regulons for predictive algorithms, we explore 4 other types of regulons and quantify their functional complexity by two different methods. Our results indicate that complex regulons, defined as a group of genes regulated by the same combination of TFs, are the most functionally homogeneous type of regulons. Finally, we measure the coexpression of genes in each type of regulon and show that, consistent with our results, complex regulons are the most coexpressed.

## MATERIALS AND METHODS

### GENSOR Unit assembly

GENSOR Units were assembled using the semi-automatic pipeline reported in (Ledezma-Tejeida, Ishida, and Collado-Vides 2017). Data from RegulonDB v9.4 (Gama-Castro et al. 2016) was obtained from two custom made datasets available at GitHub and the TF-TU and TU-genes datasets available at the RegulonDB website. All data from EcoCyc (Keseler et al. 2017) was obtained using the Pathway Tools software v19.5 (Karp et al. 2015), the Perlcyc API and custom Perl scripts. Canonical metabolic pathways and enzymatic regulatory interactions were taken from EcoCyc. Enzymatic regulatory interactions were only added to a GENSOR Unit if the regulatory metabolite and the regulated enzyme were already present in the GENSOR Unit. A modified version of the pipeline was used to assemble GENSOR Units using as input a custom group of genes.

### Connectivity

Connectivity was calculated in three steps: (1) all enzymes in the GENSOR Unit were identified (2) All metabolic fluxes in the GENSOR Unit were identified by connecting reactions that shared a metabolite, irrespective of it being a substrate or a product. (3) Enzymes participating in metabolic fluxes of more than one reaction were identified and termed “connected enzymes”. (4) The following formula was applied:

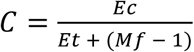

Where Ec is the number of connected enzymes (3), Et is the total of enzymes in the GENSOR Unit (1) and Mf is the number of metabolic fluxes within the GENSOR (2). Since the expectation is that all enzymes are involved in one metabolic flux, any extra metabolic fluxes are penalized in the denominator. Essentially, connectivity reflects the fraction of enzymes in the GENSOR Unit that cooperate with others in a metabolic flux (i.e. pathway), penalized by the number of unexpected fluxes. Enzymatic regulation is considered in the metric by additionally labeling as “connected enzymes” any enzyme that is regulated by a metabolite also present in the GENSOR Unit. Final calculation was performed using a custom Perl script available at GitHub.

### Regulatory groups and regulons

Regulatory groups were defined as depicted in **Table 2**. Regulons were identified through a custom Perl script using information from RegulonDB in the following manner (also see **Figure S4**):

#### General regulons

One regulon per TF. Includes all the genes that have at least one annotated binding-site for the TF in their promoter region.

#### Strict regulons

One or two regulons per TF. Includes the subset of genes in a general regulon that are regulated under the same effect (activation/repression).

#### Simple regulons

Zero or one regulons per TF. Includes all the genes that have identified binding-sites only for one TF in their promoter region.

#### Complex regulons

Zero, one or more regulons per TF. Includes all the genes that have annotated binding-sites for the same combination of TFs in their promoter region.

#### Conformation regulons

Zero, one or more regulons per TF. Includes all the genes that have annotated binding-sites in their promoter region for the same TF in complex with the same effector molecule. Only TFs in complex were considered because interactions annotated as being regulated by the TF alone are not discernable from those where no information on conformation is available.

#### Conformation + effect regulons

Zero, one or more regulons per TF. Includes the subset of genes in a conformation regulon that are regulated under the same effect (activation/repression).

Global TFs (ArcA, CRP, IHF, Fis, FNR, HNS, Lrp) were considered in all regulatory groups, except general regulons since by definition (Martínez-Antonio and Collado-Vides 2003) they are involved in more than one biological process and could introduce bias to our results. All regulons were calculated from RegulonDB datasets using custom Perl scripts.

### Metabolic Pathways, Transcription Units and Gene Ontology terms

Genes belonging to metabolic pathways were automatically retrieved using Pathway Tools v19.5. Transcription Units were obtained from RegulonDB downloadable datasets, those with only 1 gene were eliminated. Gene Ontology (GO) terms (Consortium 2000; Gene and Consortium 2017) were obtained from the PortEco Filtered Annotation File version 24/05/2017, available at the AmiGO website. Members of a GO term were expanded to include the terms that directly belong to it, plus all the genes that belong to its children terms. All analyses were performed using only the Biological Process branch of the ontology.

### Randomization of regulons

Sets of random regulons were created using a custom Perl script. For each regulatory unit, the script created a list of the genes that belonged to all of its regulons and recreated each regulon by assigning random genes from the list until the regulon reached its original gene size. Assignment of random genes allowed repetition. The process was repeated 100 times for each regulatory unit, in order to obtain 100 sets of random regulons. Random regulons for connectivity analysis were created with the same algorithm with the exception that enzyme size was maintained, as opposed to gene size. Enzymes were identified using Pathway Tools v19.5.

### Identification of dominant GO terms

We obtained the fraction of genes in a GENSOR Unit that is present in each of the 2860 Biological Process (BP) Gene Ontology (GO) terms. The GO term with the highest fraction of genes was selected as the dominant GO term.

In case of ties, the most specific term (farthest from the root) was selected. Only GENSOR Units with more than one gene annotated in the BP branch of the ontology were considered in the analysis. Genes that were not annotated in at least one term of the BP branch were excluded from the analysis. Genes that belonged to multiple GO terms were considered in all their terms.

### Coexpression of regulons

Coexpression analyses was performed using expression data from the COLOMBOS compendia (Moretto et al. 2016) across 4077 microarray contrasts. The Spearman correlation of gene expression across all conditions was computed for all possible combinations of gene pairs in a regulon. The median of the obtained Spearman correlations was calculated and used as the representing coexpression value of the regulon. Regulons with less than two genes were excluded from the analysis.

### Data availability

All regulon datasets, random regulons, GENSOR Units, custom scripts and the raw data used in this study are available at GitHub [https://github.com/dledezma/functional_homogeneity].

## RESULTS

### Assembly of GENSOR Units and calculation of connectivity

We used the GENSOR Unit framework (Ledezma-Tejeida, Ishida, and Collado-Vides 2017) to further analyze the relationship between regulons and the metabolic effect of their gene products. A general regulon is defined as the group of genes directly regulated by a TF, regardless of the effect (positive, negative or dual) of the TF and regulation from other TFs. For each TF, we automatically retrieved from RegulonDB (Gama-Castro et al. 2016) its known effectors, active and inactive conformations, regulated genes and the effect of the regulatory interactions. From EcoCyc (Keseler et al. 2017), we retrieved the gene products and any protein complex they belonged to a GENSOR Unit. If the gene products were enzymes, we extracted the catalyzed reactions, their substrates, products and directionality. Finally, we included canonical metabolic pathways in a simplified way. In each GENSOR Unit, we linked pairs of metabolites that are present in the same canonical pathway taking into account the directionality of the pathway (i.e. that one metabolite can be transformed into the other). Only one meta-reaction (called complementary pathway reaction in RegulonDB) was added to link the metabolites, regardless of the number of intermediate reactions between them in the pathway. (**Figure S1**). The end result was a multilevel network that included Transcription Units (TUs), proteins, protein complexes, and metabolites termed Genetic Sensory Response Unit or GENSOR Unit for short. GENSOR Units integrate the transcriptional and metabolic level in a single network, providing a higher-level view of their interplay and their physiological relevance (**Figure S2**).

We have previously reported (Ledezma-Tejeida, Ishida, and Collado-Vides 2017) a connectivity metric that measures the functional homogeneity of a GENSOR Unit in terms of the ability of its metabolic reactions to create a metabolic flux by sharing substrates or products, as in a pathway. In brief, connectivity takes into account the number of enzymes (Ec) whose catalyzed reactions create a metabolic flux, the total number of enzymes (Et) and the total number of metabolic fluxes (Mf) present in the GENSOR Unit (see Methods). It is calculated with the formula:

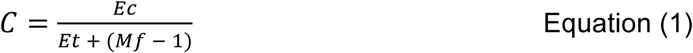

The connectivity formula returns a value from 0 to 1. Zero indicates total functional heterogeneity, where none of the reactions in the GENSOR Unit happen consecutively. A value of one indicates a paradigmatic GENSOR Unit where all the reactions are involved in the same metabolic flux and therefore, are functionally related. We calculated the connectivity of 200 GENSOR Units and eliminated those with less than two enzymatic reactions to avoid artificial values of 0. The resulting connectivity distribution (**Figure 1A**), using a different version of RegulonDB, replicates previous results (Ledezma-Tejeida, Ishida, and Collado-Vides 2017) where the largest proportion of GENSOR Units has a connectivity value of 1, followed by those with connectivity 0 and a continuum in between.

**FIGURE 1.**
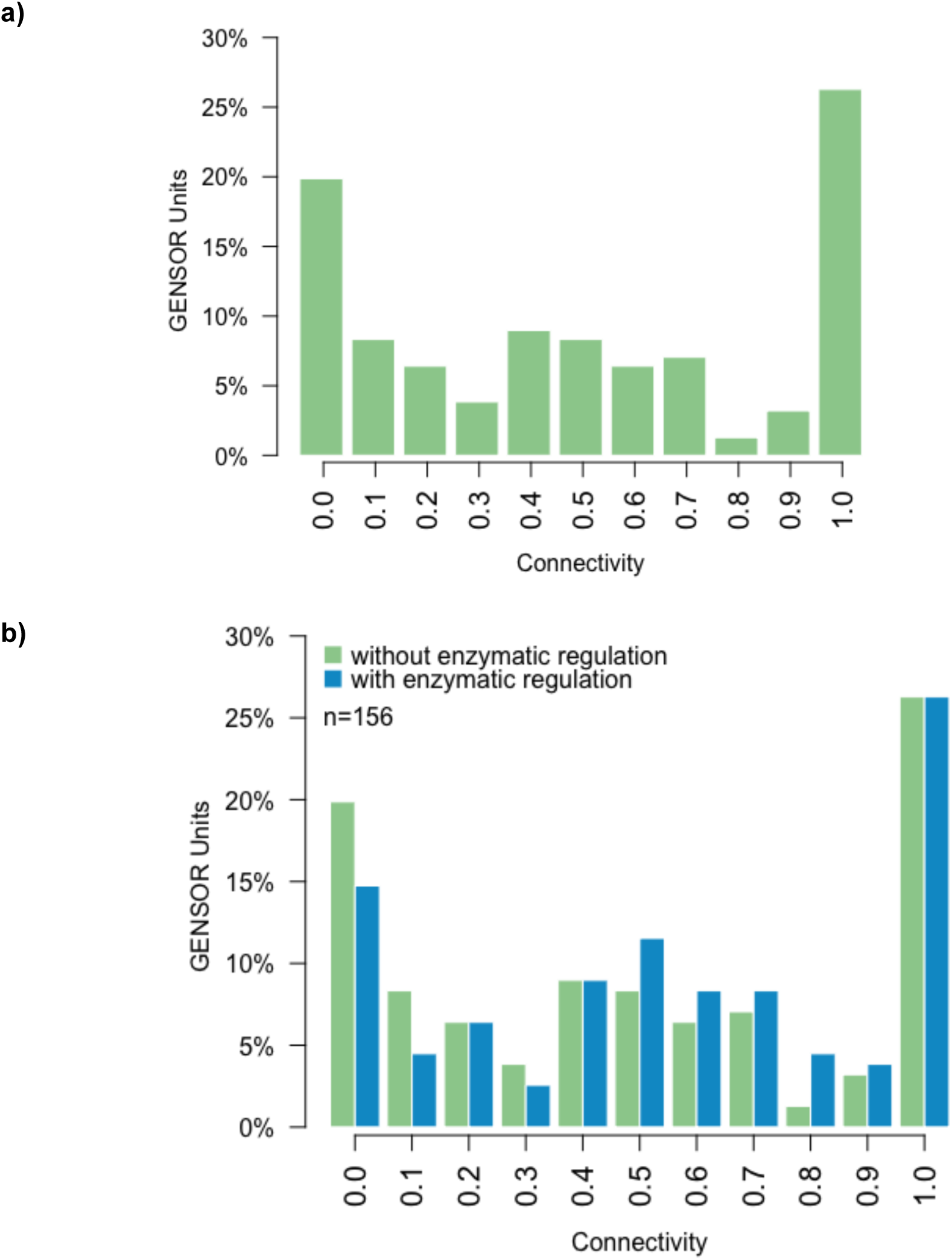
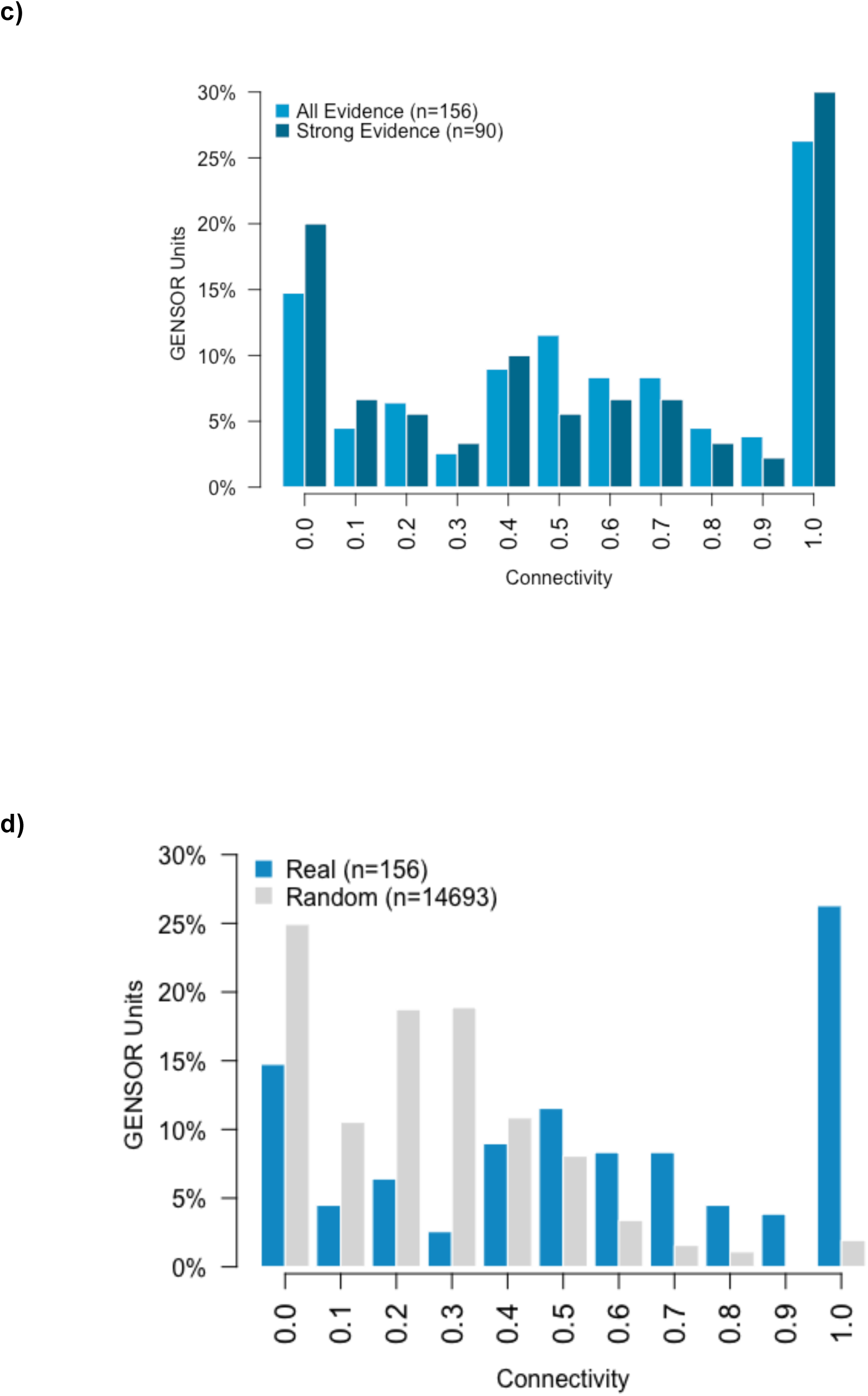
Connectivity analysis. **(a)** Connectivity distribution of GENSOR Units calculated using the previously reported algorithm on later database versions. **(b)** Distribution of modified connectivity to consider enzymatic regulation in GENSOR Units, compared to (a). **(c)** Connectivity distribution of GENSOR Units containing only regulatory interactions with strong evidence, compared to GENSOR Units including all reported regulatory interactions. **(d)** Mean connectivity distribution of GENSOR Units assembled from 100 random regulons, compared to real GENSOR Units. All distributions in (c) and (d) include enzymatic regulation.

### Functional heterogeneity in regulons is not explained by enzymatic regulation, low-confidence regulatory interactions or chance

Once we reproduced previous results, we explored artifacts that could be responsible for the observed connectivity gradient. As a first step, we examined the role of enzymatic regulation in the connectivity of GENSOR Units. It is known that enzymatic regulation plays an important role in regulatory genetic programs (Chubukov et al. 2014). For example, anthranilate synthase is the enzyme that catalyzes the first step of tryptophan biosynthesis, and it can be allosterically inhibited by L-tryptophan, the end product of the pathway (Pabst, Kuhn, and Somerville 1973). This type of interactions can create functional links between enzymes and metabolites that are missed in the connectivity metric but are important because they increase the functional homogeneity of GENSOR Units. We retrieved 398 enzyme-metabolite interactions annotated in EcoCyc and were included in 80 GENSOR Units. The connectivity calculation was slightly modified by expanding the definition of a *connected enzyme* (Ec in equation 1), from “*an enzyme whose catalyzed reaction shares at least one substrate or product with another reaction in the GENSOR Unit*” to “*an enzyme whose catalyzed reaction shares at least one substrate or product with another reaction in the GENSOR Unit or is allosterically regulated by the substrate or product of the reaction catalyzed by another enzyme in the GENSOR Unit*”. The resulting connectivity distribution using the modified connectivity (**Figure 1B**) does not differ significantly from the original distribution (Wilcoxon-Mann-Whitney test; p-value > 0.05). This result suggests that the gradient of functional complexity originally observed is not a consequence of missing metabolite-enzyme interactions. Nevertheless, metabolite-enzyme interactions add useful information to GENSOR Units, so we included them and used the modified version of connectivity in all further analyses.

The gradient of connectivity could also be explained by spurious regulatory interactions. RegulonDB classifies each regulatory interaction in strong or weak according to the evidence provided by the methods used to identify it (Weiss et al. 2013). For instance, binding experiments of the purified TF are considered strong evidence, while changes in gene expression in a TF mutant strain are considered weak evidence. Certain combinations of multiple independent weak evidence can also add up for a strong evidence. It is possible that interactions with low levels of confidence such as those inferred from a mutant phenotype (i.e. the TF mutant strain shows a phenotype in which the regulated gene is involved) do not happen in reality and are introducing noise in the connectivity of the GENSOR Unit that includes them. We used RegulonDB evidence codes to eliminate all the regulatory interactions with weak evidence and assembled high-confidence GENSOR Units. If the connectivity gradient observed so far is being biased by low-confidence interactions, the connectivity distribution of high-confidence GENSOR Units should differ. **Figure 1C** shows that this is not the case since the distribution is not significantly different (Wilcoxon-Mann-Whitney; p-value > 0.05), even considering that the total GENSOR Units tested decreased by 43% and the average size of the assembled GENSOR Units decreased from 5.6 enzymes to 3.0. As we have shown before (Ledezma-Tejeida, Ishida, and Collado-Vides 2017), there was no correlation between the connectivity value of a GENSOR Unit and its size. This result shows that low-confidence interactions are not introducing bias to our results, and are not responsible for the connectivity gradient.

The prevalence of the connectivity gradient led us to explore whether it could be expected by chance, meaning that we are not measuring a biological property, but the result of a metric’s artifact. In order to do this, we took the complete set of genes in the Transcriptional Regulatory Network (TRN) and recreated the 200 regulons by randomly selecting genes from the set. The only property of the regulons that remained was the number of enzymes since, as mentioned before, the connectivity value does not depend on the GENSOR Unit size. We assembled GENSOR Units using the random regulons as starting points and obtained their connectivity distribution. For statistical significance we repeated this process 100 times and obtained the mean connectivity distribution (**Figure 1D**). The resulting distribution is significantly different from the original connectivity distribution (Wilcoxon-Mann-Whitney; p-value < 2.2e-16), suggesting that the connectivity gradient observed in regulons is not an artifact. The three previous results support that the gradient of connectivity reflects underlying biological principles, reinforcing the notion that functional homogeneity is not a general property of individual TFs.

### The Gene Ontology also reveals functional heterogeneity in regulons

So far we have supported the functional heterogeneity of GENSOR Units in terms of the connectivity metric, next, we explored whether other methods of functional quantification yield the same results. Our connectivity metric reflects the functional homogeneity of a GENSOR Unit under the assumption that proximity in the metabolic network implies similar function, as in a metabolic pathway. This assumption is commonly accepted since enzymes and metabolites that are present in the same metabolic pathway (e.g. KEGG or EcoCyc pathways), and therefore are close in the larger metabolic network (Feist et al. 2007), work together to produce a final product. Nevertheless, there are other approaches to describe functional homogeneity, of relevance are those based on the Gene Ontology (GO) (Consortium 2000; Gene and Consortium 2017), a functional classification of genes mainly based on homology and phenotypic effects of genes. One of the most used approaches is the GO Enrichment Analysis, which relies on statistical analysis to identify functional terms that are over or under-represented in a set of genes of interest, given the background of the functional annotations of each gene in the genome (Yon Rhee et al. 2008; Mi et al. 2017). Essentially, the GO Enrichment Analysis answers the question: “Which biological processes are significantly enriched in a GENSOR Unit?”. Unfortunately, the results are not a true reflection of functional homogeneity since genes tend to belong to more than one GO term and different subsets of genes in a GENSOR Unit could account for different enriched processes, giving no direct information on whether there is one process where all the genes in the GENSOR Unit are working together. For a better reflection of functional homogeneity we focused on two different questions: “How general is the biological process that can simultaneously describe all the genes in a GENSOR Unit?”, and “What is the fraction of a GENSOR Unit that can be explicitly explained by a biological process?”.

To answer the first question, we identified the GO term that describes all the genes in each GENSOR Unit and focused on the tree structure of the ontology to quantify how specific the term is. We obtained, for each GENSOR Unit, the subset *G* of genes that are present in the Biological Process (BP) branch of GO, eliminating those GENSOR Units with less than two annotated genes to avoid uninformative results. For the remaining 185 GENSOR Units we identified the farthest downstream GO term from the root of the ontology that included all the genes in the GENSOR Unit, in other words, the biological process that is most representative of, or dominant, in the GENSOR Unit. Since terms closer to the root are more general in their definition, picking the term farther from it implies that it will also be the most informative possible. The root term, “Biological Process” was labeled as level 1, its immediate children as level 2 and so on until level 11. The worst case scenario would be a GENSOR Unit whose most representative GO term is in level 1 of the ontology, indicating that no other term in the ontology can fully describe that GENSOR Unit. This scenario would also imply functional heterogeneity given that the genes have functions spread across the ontology. To validate this interpretation, we obtained the dominant GO term of global regulators (ArcA, CRP, Fis, FNR, HNS, IHF, Lrp), which by definition are involved in several biological processes. The dominant GO term of all global regulators is indeed level 1 term “Biological Process”, the most general term. Results for the complete set of GENSOR Units show that 68.1% also have level 1 term “Biological Process” as dominant GO term **(Figure 2A)**. Additionally, 9.1% of GENSOR Units had a level 2 GO term as its best descriptor. Taken together, these results agree with connectivity results in that around three quarters of GENSOR Units are functionally heterogeneous. The high percentage of functionally heterogeneous GENSOR Units is not a consequence of the ontology having more terms in levels 1 and 2, in fact these levels only include 20 terms, accounting for 0.005% of total BP GO terms (**Figure S3A**). Another explanation could be that the more general terms are the only ones with enough genes annotated to describe all the genes in a GENSOR Unit, but that is also not the case since 124 GO terms of level 3 and higher contain more genes than the largest GENSOR Unit (**Figures S3B-C**). A possible functional bias is that TFs are annotated in the ontology with terms related to transcription, as opposed to the processes they regulate. To explore this possibility, we excluded autoregulated TFs from the analysis. Results were not significantly affected since 69.7% of GENSOR Units still obtained a dominant GO term in levels 1 or 2. It is also possible that expecting a complete regulon to be explained by a single GO term is extremely stringent and maybe two GO terms are enough to explain all regulons. However, that is not the case since allowing two dominant GO terms only increases by 30.8% our interpretative power, leaving 46.4% of GENSOR Units still dispersed in three or more biological processes. This analysis supports the notion that functional heterogeneity is a general property of regulons.

**FIGURE 2.**
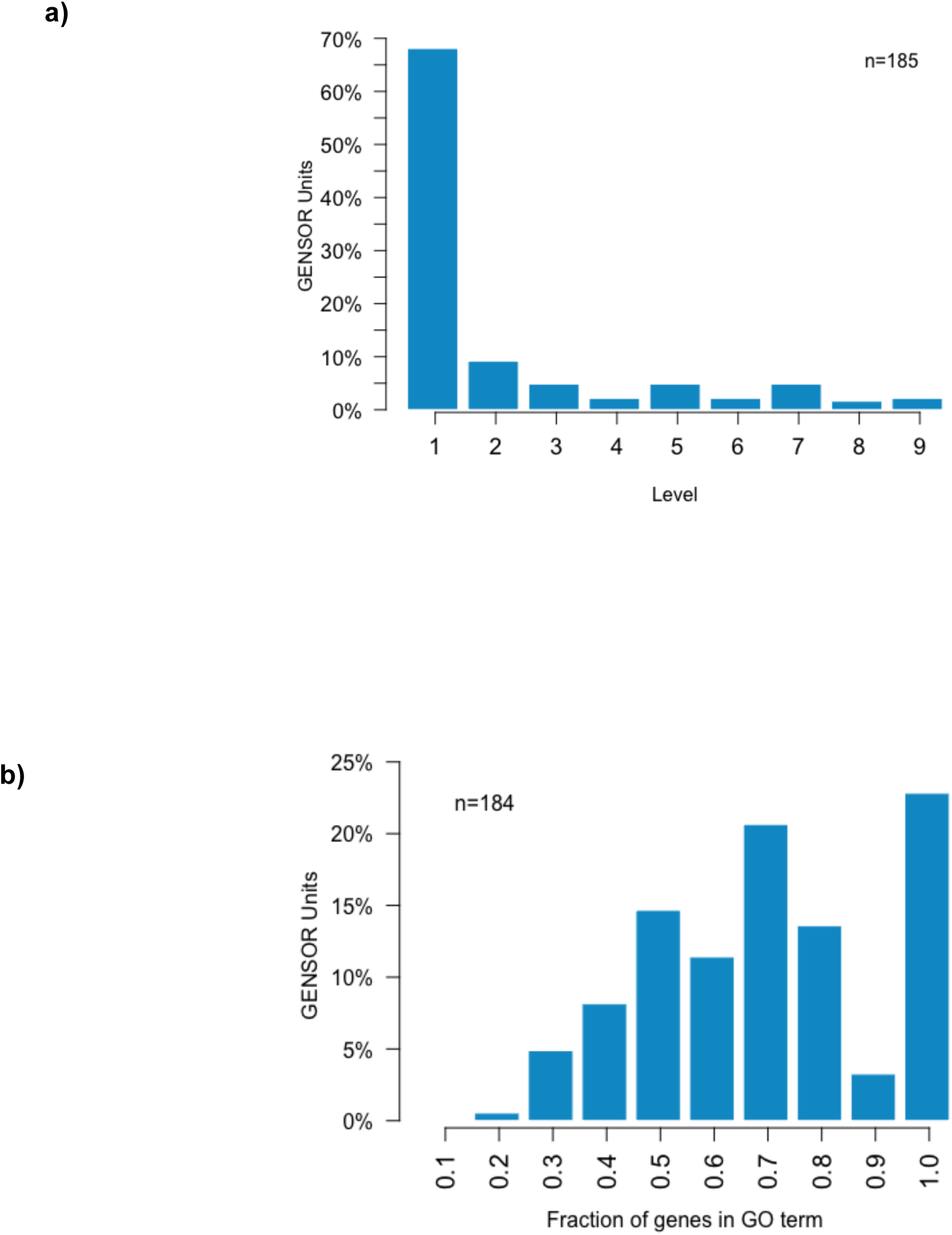
GO analysis. **(a)** Distribution of levels of the GO term that best describes each GENSOR Unit. **(b)** Distribution of the highest fraction of genes of each GENSOR Unit that are present in the same GO term. Only descriptive GO terms (level 3 and higher) were considered in this analysis. In both analyses, GENSOR Units with less than 2 genes annotated in the ontology were excluded.

The second question, “What is the highest fraction of a GENSOR Unit that can be explicitly explained by a biological process?” was answered by eliminating the two most general levels of the ontology, and repeating the analysis. By eliminating the most general processes we are identifying the most representative GO term of each GENSOR Unit that is still informative. We obtained the fraction of genes in the GENSOR Unit that are included in the best GO term, the closer the fraction is to 1, the more functionally homogeneous the GENSOR Unit. The best scenario will be GENSOR Units with a fraction of 1, meaning that all of their genes are involved in an informative Biological Process. **Figure 2B** shows that the gradient of complexity observed in connectivity analysis is also present here: most GENSOR Units cannot be entirely described through a single, informative Biological Process, and the fraction of genes that belong to the most representative process varies from 0.2 to 1. Notably, level 6 is the most populated in the ontology (**Figure S3A**) with 25.5% of all terms belonging to it, but only 10.3% of GENSOR Units have their most representative GO term at that level (**Figure S3D**), suggesting that the analysis was not biased by properties of the ontology. Level 3, the most general in this analysis, was the most represented level (36.4%) in GENSOR Units (**Figure S3D**), further supporting their functional heterogeneity.

Functionally heterogeneous GENSOR Units are not the less studied TFs. For example, MetJ is a TF associated with biosynthesis and transport of methionine (hence its name). As expected, its regulated genes are included in methionine-related GO terms such as “methionine biosynthetic process” or “protein methylation”, but they only include a low fraction of the regulon. Interestingly, MetJ’s dominant GO term is “organonitrogen compound metabolic process”, which includes 58% of the regulated genes and is not directly related to methionine.

As mentioned before, connectivity and GO analyses rely on different properties of functional relationships, but both suggest that only around a quarter of TFs regulate functionally homogeneous genes. Scores do not show any clear correlation between the metrics (**Figure S3E**), but they overlap in 15 GENSOR Units that scored perfect functional homogeneity in both analyses: AccB, AllS, ArsR, BetI, BirA, CynR, FabR, FeaR, GcvA, MazE, MazE-MazF, MhpR, TreR, XapR, and YqjI. The dominant GO term of these regulons and a more detailed description of their GENSOR Units can be found in **Table 1**. Effectors are known for 10 of them. In most cases, the annotated biological process is directly related to the effector, for example, AllS is involved in “allantoin assimilation pathway” and it binds to allantoin (Rintoul et al. 2002). In other cases, the relationship was more indirect but still present, for instance BetI’s effector is choline (Rkenes, Lamark, and Strøm 1996) and is annotated as being involved in “response to stress” because choline can be converted into glycine betaine, a thermo- and osmoprotectant (Caldas et al. 1999) by genes directly regulated by BetI. These GENSOR Units reflect the most functionally local TFs: involved in a single biological process and placed at the bottom of the TRN hierarchy since they do not regulate other TFs.

**TABLE 1.** Functionally homogenous GENSOR Units. Regulated biological processes shown are obtained from the most representative GO term identified. Summaries describe active conformations and end/metabolites of the metabolic fluxes mediated by the TF of the GENSOR Unit.

| **GENSOR Unit** | **Known Effector** | **Dominant GO term (level of term)** | **Mechanistic summary of GENSOR Unit** |
| --- | --- | --- | --- |
| AccB |  | monocarboxylic acid biosynthetic process (7) | AccB, by itself, negatively regulates the expression of genes required to produce malonyl-CoA. |
| AllS | allantoin | allantoin assimilation pathway (7) | AllS, in the presence of allantoin, positively regulates the expression of genes required to produce ammonium and oxalurate. |
| ArsR | arsenite<br>antimonite | response to arsenic-containing substance (4) | ArsR, by itself, negatively regulates the expression of genes required to produce oxidized glutaredoxin and arsenite. |
| BetI | choline | response to stress (3) | BetI, by itself, negatively regulates the expression of genes required to produce betaine and A(H<sub>2</sub>). |
| BirA | biotinyl-5'-AMP | monocarboxylic acid biosynthetic process (7) | BirA, in the presence of biotinyl-5'-AMP, negatively regulates the expression of genes required to produce coenzyme A, 5'-deoxyadenosine, AcpP, adenosylhomocysteine, S-adenosyl-4-methylthio-2-oxobutanoate, oxidized [2Fe-2S] ferredoxin, methionine, malonyl-[acp] methyl ester, unsulfurated [sulfur donor] and biotin. |
| CynR | cyanate | cyanate catabolic process (6) | CynR, in the presence of cyanate, positively regulates the expression of genes required to produce carbamate. |
| FabR | | monocarboxylic acid<br>monocarboxylic<br>monobiosynthetic process<br>(7) | FabR, by itself, negatively regulates the expression of genes required to produce 3-oxo-decanoyl-[acp], 3-oxo-dodecanoyl-[acp], trans tetradec-2-enoyl-[acp], trans hexadecenoyl-[acp], (2E)-dodec-2-enoyl-[acp], 3-ketopimelyl-[acp] methyl ester, 3-oxo-cis-delta7-tetradecenoyl-[acp], 3-oxo-cis-delta9-hexadecenoyl-[acp], 3-oxo-octanoyl-[acp], 3-oxo-palmitoyl-[acp], 3-oxo-hexanoyl-[acp], 3-oxo-myristoyl-[acp], trans hex-2-enoyl-[acp], $\text{CH}_3(\text{CH}_2)_x\text{CH}=\text{CHCO-S-ACP}$, crotonyl-[acp], acetoacyl-ACP and acetoacetyl-[acp] |
| FeaR | | cellular biogenic amine<br>catabolic process (7) | FeaR, by itself, positively regulates the expression of genes required to produce perhydrol, oxopropanal, $\text{RCHO}$, ammonium and phenylacetate. |
| GcvA | purine<br>glycine | nitrogen compound<br>metabolic process (3) | GcvA, by itself, dually regulates the expression of genes required to produce [glycine-cleavage complex H protein] N6-dihydrolipoyl-L-lysine, methylene-H4PteGlu(n) and ammonium. |
| MazE |  | nucleobase-containing<br>compound catabolic<br>process (5) | MazE, by itself, negatively regulates the expression of genes required to produce cytidylate, dAMP, 5'-IMP, 5'-UMP, dGMP, dCMP, dTMP, xanthosine-5-P, dIMP, mononucleotide and dUMP. |
| MazE-MazF |  | nucleobase-containing<br>compound catabolic<br>process (5) | MazE-MazF, by itself, negatively regulates the expression of genes required to produce cytidylate, 5'-IMP, |
|  |  |  | 5'-UMP, dCMP, dAMP, dIMP, dGMP, xanthosine-5-P, mononucleotide, dTMP and dUMP. |
| MhpR | 3-(2,3-dihydroxyphenyl)propanoate 3-(3-hydroxyphenyl)propanoate | aromatic compound catabolic process (5) | MhpR, in the presence of 2,3-DHP, positively regulates the expression of genes required to produce succinate, (2Z)-2-hydroxypenta-2,4-dienoate, acetyl-CoA, fumarate and pyruvate<br>MhpR, in the presence of 3HPP, positively regulates the expression of genes required to produce succinate, (2Z)-2-hydroxypenta-2,4-dienoate, acetyl-CoA, fumarate and pyruvate |
| TreR | trehalose 6-phosphate trehalose | cellular metabolic process (3) | TreR, by itself, negatively regulates the expression of genes required to produce glucose-6-P, PtsH and glucopyranose. |
| XapR | xanthosine | nucleobase-containing small molecule metabolic process (4) | XapR, in the presence of xanthosine, positively regulates the expression of genes required to produce xanthine, hypoxanthine, guanine and nicotinamide ribose.<br>XapR, by itself, positively regulates the expression of genes required to produce xanthine, hypoxanthine, guanine and nicotinamide ribose. |
| YqjI | nickel Fe+2 | cellular response to stimulus (3) | YqjI, by itself, negatively regulates the expression of genes required to produce (2,3-dihydroxybenzoylserine)3, Fe+2 and siderophore. |

### Complex regulons are the most functionally homogeneous type of regulons

Having further supported the notion that the genes directly regulated by a TF are not generally involved in the same biological process, we focused on exploring the functional homogeneity of other types of regulons, defined slightly different. Our underlying assumption is that transcriptional regulation plays a central role in cellular decision-making: when a cell is faced with a change in the environment, a coherent response must be orchestrated mainly through the action of TFs. The resulting hypothesis is that there should be a type of regulatory unit in the TRN that also acts as a functional unit. We selected 3 previously reported (Gutierrez-Rios et al. 2003) types of regulatory groups: simple, complex and strict regulons (**Table 2**). Their definitions (**Table 2, Table S1, Figure S4**) combine two properties of TFs: their effect on genes, and their shared occupation of promoters with other TFs. Based on previous reports on the relevance of TF conformation information (Gutierrez-Rios et al. 2003); we also considered the groups of genes directly regulated by a specific TF-effector complex, and those directly regulated by a specific TF-effector complex under the same effect. For instance, the genes activated by the complex TyrR-phenylalanine belong to a different regulon than those repressed by TyrR-tyrosine. In total, we analyzed 6 types of regulatory groups (**Table 2, Table S1, Figure S4**). We obtained the groups of genes in the TRN derived from each definition and used them as starting point to assemble GENSOR Units. As a positive control we also used the GENSOR Unit assembly pipeline on groups of genes defined by pathways and GO terms. As negative control, we generated 100 sets of random gene groups for each type of regulatory grouping (see Methods). To compare the functional homogeneity of regulatory groups, we obtained their connectivity distribution and identified dominant GO terms as described in the previous sections.

**TABLE 2.** Definitions of regulatory units and controls used to assemble GENSOR Units.

| **Regulatory Groups/Controls** | **Definition** | **Total regulons** |
| --- | --- | --- |
| General Regulons | Genes directly regulated by a TF. Irrespective of effect, TF conformation, or coregulation with other TFs. | 200 |
| Strict Regulons | Genes directly regulated by a TF, under the same effect (+/-), irrespective of TF conformation, or coregulation with other TFs. | 278 |
| Simple Regulons | Genes directly regulated one and only one TF. Irrespective of effect or TF conformation. | 106 |
| Complex Regulons | Genes directly regulated by a combination of TFs, irrespective of effect or TF conformation. | 395 |
| Conformation Regulons | Genes directly regulated by a specific conformation (TF-effector complex) of a TF, irrespective of effect, or coregulation with other TFs. | 87 |
| Conformation + effect regulons | Genes directly regulated by a specific conformation (TF-effector complex) of a TF, under the same effect (+/-), or coregulation with other TFs. | 132 |
| GO term | Genes annotated in the same GO term, plus all the genes on its children terms | 2860 |
| Pathway | Genes that belong to the same metabolic pathway | 420 |
| Transcription Unit | Genes transcribed on the same mRNA molecule | 1026 |

Connectivity values (**Figure 3A** and **Table 2**) show that the highest scoring regulatory group, with a median of 0.8, are complex regulons - groups of genes regulated by the same combination of TFs. The rest of regulatory groups have a median connectivity of 0.6, except for simple regulons with a lower value of 0.5 (**Table S2**). Pathways, the positive control, have a median of 1, as expected. All regulatory groups have a distribution significantly different from random values (**Figures S5A-G**). Consistent with connectivity results, GO analysis also scored complex regulons as the most functionally homogeneous regulons with a median fraction of genes of 1.0 (**Figure 3B**). The rest of regulons showed a median of around 0.7, with the lowest being general regulons on 0.67 (**Table S2**).. The positive control, groups of genes belonging to a GO term, has only 1.0 values, as expected. Again, all regulatory groups have a distribution significantly different from random values (**Figures S6A-G**). Through these two independent analyses we have shown that complex regulons are the most functionally homogeneous regulatory group from the ones tested, which fits with a model of TF coordination to orchestrate a cellular decision. Results suggest that functionality in the TRN should be understood in terms of the cooperation between TFs and not at the level of individual TFs.

**FIGURE 3.**
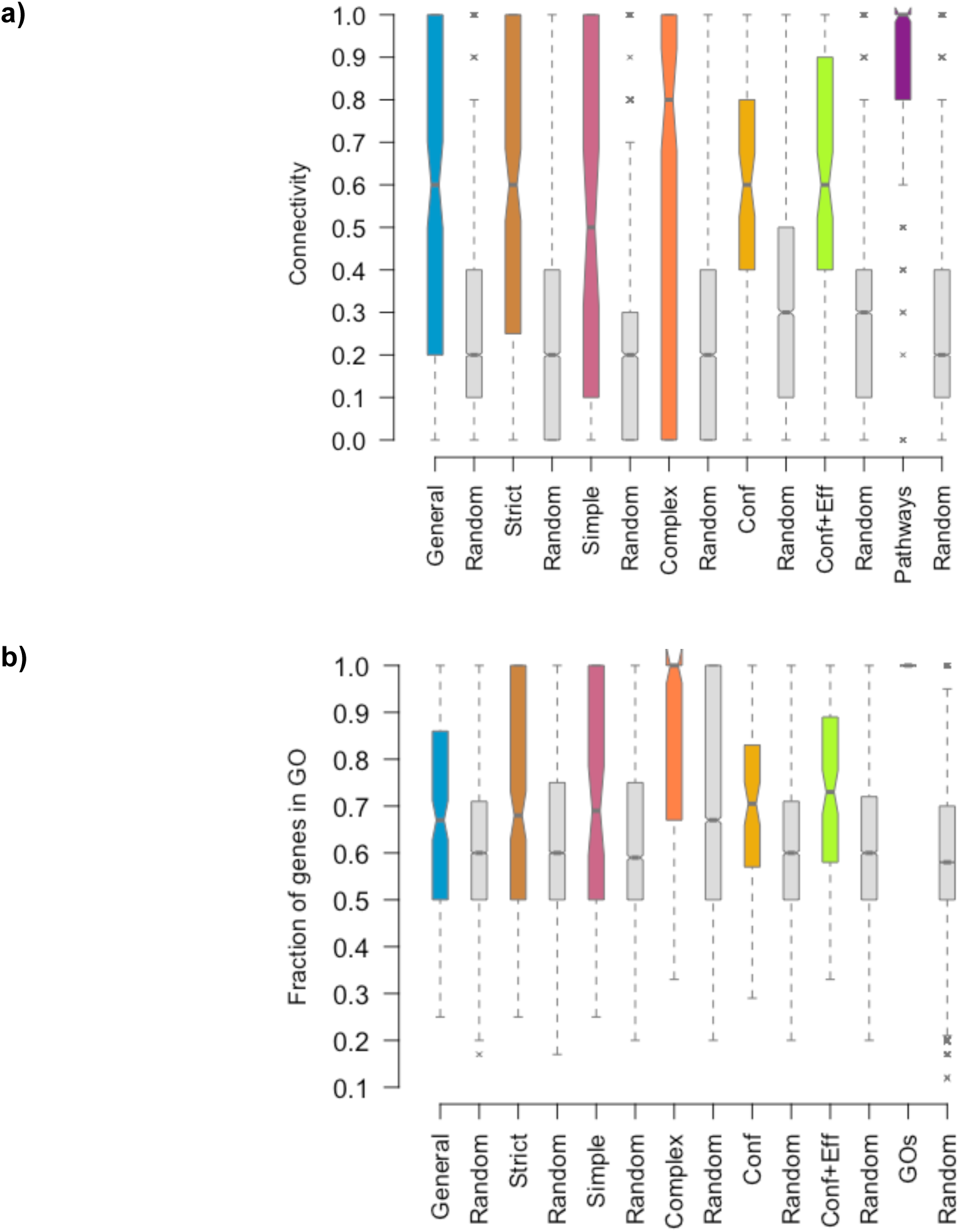
Functional homogeneity of different regulatory units. **(a)** Boxplots of connectivity distributions of regulatory groups and 100 randomizations of each set of regulons. Pathways are shown as control. **(b)** Boxplots of distributions of the highest fraction of genes of each GENSOR Unit that are present in the same GO term, for each type of regulatory group and 100 randomizations of the genes that belong to each regulon. GO terms are shown as control. *Conf* stands for Conformation Regulons and *Conf+Eff* stands for Conformation + effect regulons. Regulon definitions can be found on table 2.

### Gene expression data supports functional homogeneity of complex regulons

To place our results on a more biologically relevant context, we quantified the coexpression of each regulon of each type of regulatory unit. Our underlying assumption is that functionally related genes should be more coexpressed than random genes, given that the execution of a biological process requires the presence of all the genes involved in it. Therefore, complex regulons, as the most functionally homogeneous type of regulatory unit, should also have the most coexpressed regulons. We measured coexpression by using COLOMBOS database (Moretto et al. 2016), a compendia of microarray experiments that include expression data for 4321 genes across 4077 contrast conditions. For each regulon, we selected all the possible gene pairs and calculated the Spearman correlation of their expression values across all conditions. The median of the correlations of all gene pairs in a regulon was used as the coexpression score of the complete regulon. Spearman correlation has been previously reported as the best statistic for coexpression analysis (Pannier et al. 2017). As a control, we included Transcription Units, genes that are transcribed on the same mRNA molecule and therefore coexpressed. Results show that most regulatory units have a median coexpression correlation coefficient of around 0.2 (**Figure 4A**), except for complex regulons, with a median of 0.39, higher than Transcription Units’ median of 0.37. Given that correlation coefficients are not very high, we cannot make inferences about coexpression properties of regulatory units. However, we have obtained evidence that complex regulons are the most coexpressed regulatory unit, as coexpressed as Transcription Units, which agrees with their observed functional homogeneity.

**FIGURE 4.**
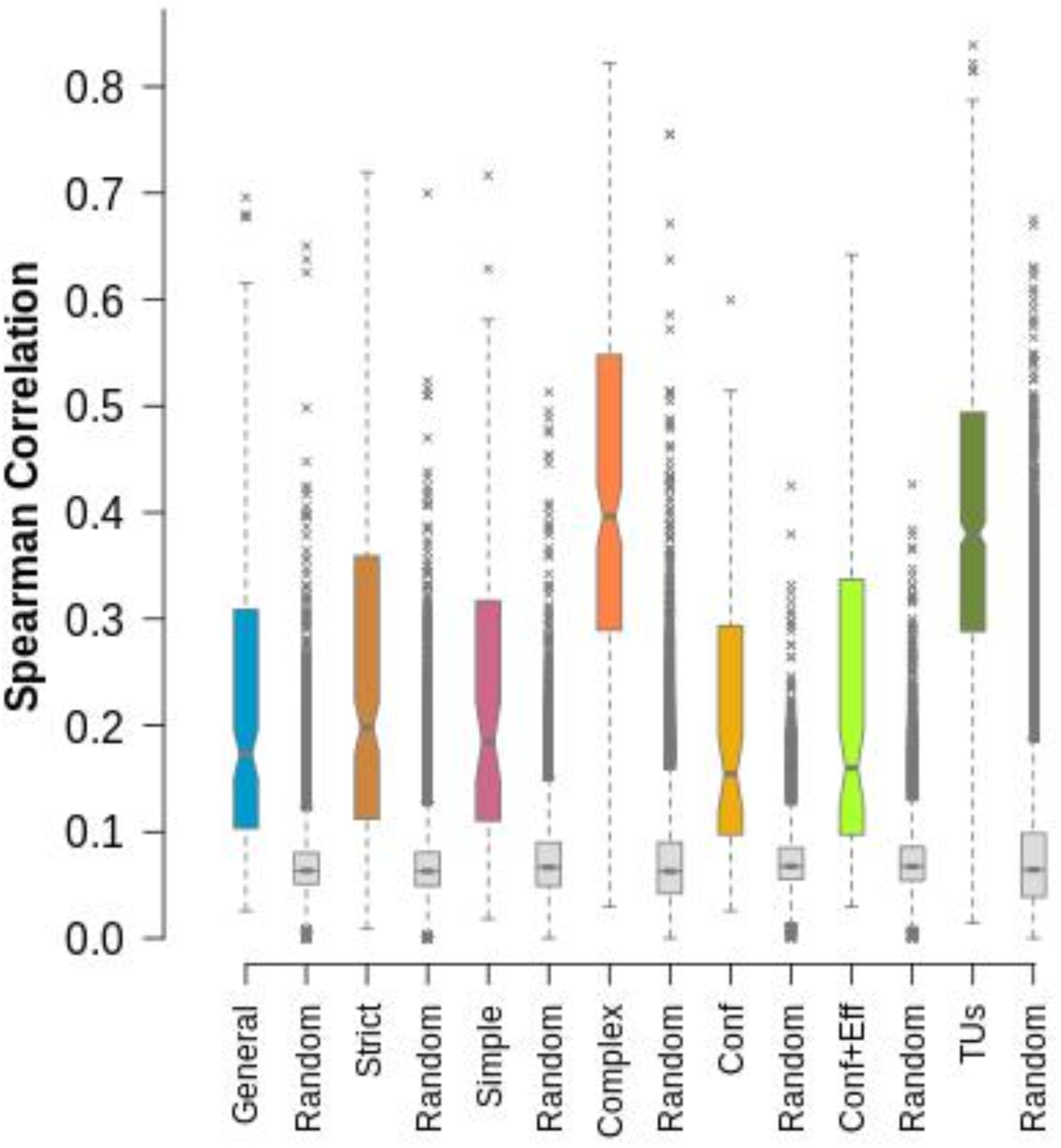
Boxplots of the distribution of coexpression values for each regulatory unit, compared to their random sets. Transcription units are shown as control.

## DISCUSSION

The common model of transcriptional regulation mediated by TFs states that an individual TF detects a specific signal, changes conformation and activates/represses a fixed set of genes. In turn, the regulated genes jointly orchestrate a response to the presence of the initial signal. Genomic studies allow the reevaluation of models, identifying the true general principles. We have shown before (Ledezma-Tejeida, Ishida, and Collado-Vides 2017) that feedback between signals and the orchestrated response is a common occurrence. On this work, we expand our analysis on the functional homogeneity of regulons by considering enzymatic regulation, the quality of data annotations, expectations by chance, and other definitions of functionality and regulons. All results confirm that only one quarter of known TFs regulate functionally homogeneous genes, evidencing that genes directly regulated by a TF are not generally involved in the same biological metabolic process. Additionally, we have shown that complex regulons are the most homogeneous regulatory unit in the TRN, which is supported by being also the most coexpressed.

The functional heterogeneity measured here assumes that all the regulatory interactions in a regulon happen simultaneously, mainly due to the lack of information on growth conditions where each regulatory interaction is active. It is possible that once this information becomes available, functional homogeneity of condition-dependent regulons will increase. However, there are reports of ChIP-seq (Fitzgerald, Bonocora, and Wade 2014), ChIP-exo (Seo, Kim, Szubin, et al. 2015; Seo, Kim, O’Brien, et al. 2015) and microarray (Göhler et al. 2011) experiments where regulated genes are involved in functions unrelated to the phenotype being studied, suggesting that even under very specific conditions our conclusions stand - direct targets of an individual TF are not necessarily involved in the same process. The most practical implication of our results is that coregulation is a dangerous assumption to propagate functional annotations to less studied genes, given that this will be correct only in around 25% of the instances, as has been shown in this analysis of a comprehensive collection of regulons in *E. coli* (**Figures 1B, 2B**). It is certainly difficult to believe that our observations are unique to the biology of *E. coli*. Confidence that coregulation implies shared function can increase to around 50% if annotations are propagated to genes regulated by the same combination of TFs (**Figures S5C, S6C**). TFs are commonly classified in local and global; one of the common features of global regulators is that the gene products they directly regulate are involved in several functional classes. Results presented here show that being involved in many functional classes is also a property of local TFs, suggesting that this criterion should not be used alone for the classification of newly discovered TFs.

On a wider perspective, we consider there are three main possible explanations for the functional heterogeneity observed in general regulons: (1) we did not evaluate the regulatory groups that drives defined biological processes; (2) there is not yet a functional framework that can describe the relationship between a regulon and its physiological effects; (3) regulation of gene expression does not rely only on TFs. The first explanation relies on the fact that we only tested 6 regulon definitions. Although we considered several mechanistic properties of TFs in these definitions, it is possible that there is a definition of regulon that has perfect correspondence with known biological processes, but we have yet to identify it.

The second explanation, that our current functional classifications of genes are not able to capture the functional regulatory logic, relies on the notion that regulation drives cellular decision-making. So far, function has been studied from the perspective of biochemical properties, homology and phenotypic effects of mutants (Consortium 2000; M. Riley et al. 2006; Galperin et al. 2015), not always taking into account regulation. Although in this work we do not comprehensively consider all available functional annotations, other widely used classifications such as COG functional terms (Galperin et al. 2015) and MultiFun terms (Monica Riley 1993) rely on similar evidences from those used by the Gene Ontology Consortium, and we chose the latter given its higher level of curation and maintenance efforts. It is noteworthy that in the original MultiFun publication (Monica Riley 1993) it is mentioned that regulation was not considered in the gene classification because even operons were not always related in metabolic terms. There could be an unexplored level of functional complexity where TFs are the main drivers, and more work would be needed to explain how apparent different processes are in fact part of a larger response to a specific signal. Examples of this interpretation efforts exist (Aquino et al. 2017; Gao et al. 2018), and they evidence our lack of a standardized functional vocabulary to interpret regulation, for instance, each ChIP-seq experiment requires a new effort by an expert curator. In the Jacob/Monod paradigm it is counter-intuitive that two genes regulated by the same TF are not be involved in a common function. It may well be that the results shown here are more due to the current limitations in gene annotations, and that the heterogeneity observed is showing our ignorance of a regulatory logic waiting to be discovered.

The third explanation, that TFs are not the main drivers of cellular decision-making, is more feasible when one places TFs in the wider context of the cell. Bacteria do not rely solely on TFs to regulate their physiology, they also depend on ribosome abundance (Borkowski et al. 2016), RNA polymerase availability (Klumpp and Hwa 2008) and intrinsic stochasticity (Munsky, Neuert, and van Oudenaarden 2012), not to mention post-translational and post-transcriptional modifications. There are estimates for the TF Cra, that it only accounts for 32% of changes in gene expression of target genes in central metabolism (Kochanowski et al. 2017). In this bigger picture, individual TFs are just one of the players in cellular decision-making, a resource to fine-tune the expression of genes that are not necessarily involved in the same process.

The observation that complex regulons are the most homogeneous regulatory unit agrees with the third explanation, since it supports that genetic programs are encoded beyond individual TFs. In this scenario, general regulons show the regulatory potential of TFs, but the specific subset of genes in the regulon that is expressed at a certain time is defined by the combinatory logic of the TFs bound to each gene’s promoter. The possible combinations of TFs, promoters, number of sites, TF effects and order of TF binding far exceed the possible biological processes regulated by a simple model of one signal - one TF - one response (Buchler, Gerland, and Hwa 2003; Mayo et al. 2006; Hunziker et al. 2010; Ezer, Zabet, and Adryan 2014; Semsey 2014). It has been shown that the large number of possible combinations allows for a faster evolution of regulatory networks (Mayo et al. 2006), which is known to happen (Borneman et al. 2007; Shou et al. 2011). The high functional homogeneity of complex regulons highlights the importance of further exploring the combinatory logic of TFs on promoters. Although the TRN of *Escherichia coli* has been widely studied, we are just beginning to explore the complexity of its relationship to metabolism and physiology. Dissecting the molecular decision-making processes associated to changes of growth conditions at a genomic level is doable with current technologies, and will no doubt bring an invaluable resource to further expand our understanding of microbial cell biology.

## Supporting information

## ACKNOWLEDGEMENTS

Authors wish to thank Luis José Muñiz-Rascado, Heladia Salgado and César Bonavides-Martínez for their skillful technical support. Mishael Sánchez-Pérez, Laura Gómez-Romero, Socorro Gama-Castro and Elad Noor for insightful discussions.

