## Supplementary material for "Limits to a classic paradigm: Most transcription factors regulate genes in multiple biological processes"

a)

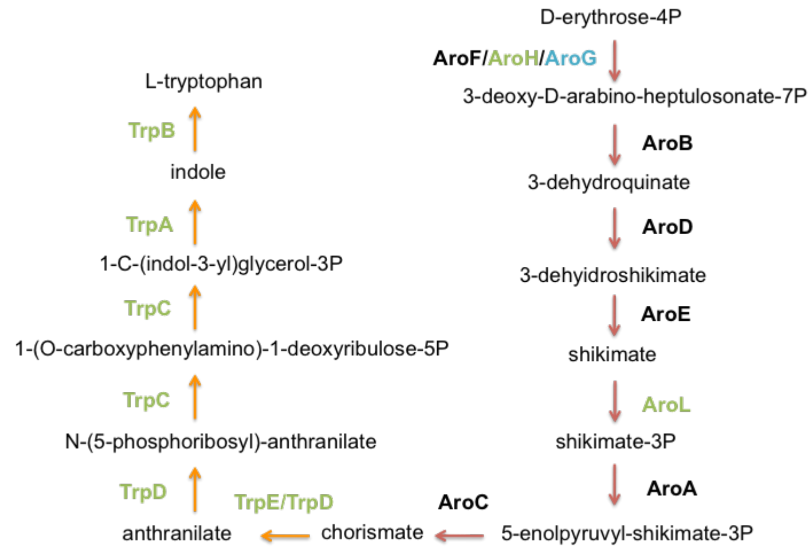


b)

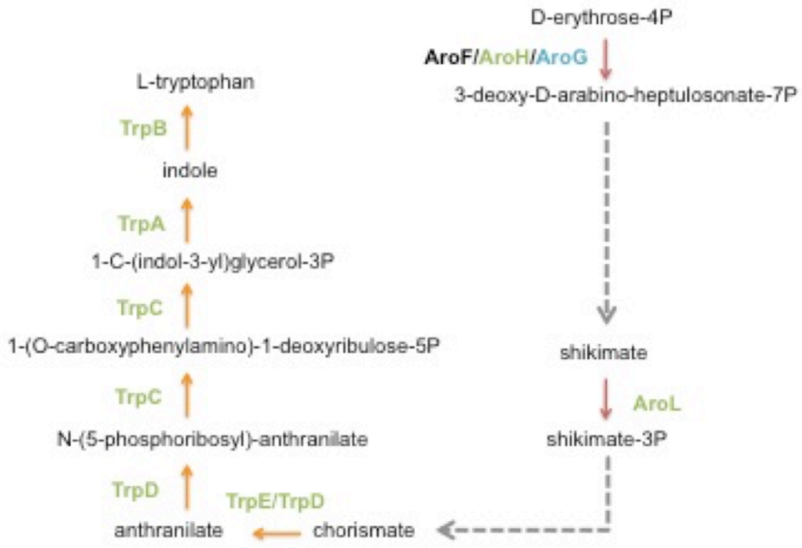


**Figure S1.** Addition of canonical metabolic pathways. Arrows depict Enzymatic reactions. Enzymes that catalyse the reaction are shown on the side of the arrow. **(a)** TrpR regulates 7 enzymes involved in the production of L-tryptophan from D-erythrose-4P (shown in green). Considering only TrpR direct targets it would appear that the reactions catalysed by AroH and AroL are not related, however the metabolic pathways of chorismate biosynthesis from 3-dehydroquinate and 3-dehydroquinate biosynthesis I indicate the existence of a metabolic flux that converts 3-deoxy-D-arabino-heptulosonate-7P into shikimate and shikimate-3P into chorismate respectively. **(b)** In the GENSOR Unit the reactions catalysed by AroB, AroD and AroE are summarized in one reaction without indication of the involved enzymes or intermediate metabolites. The same applies for reactions catalysed by AroA and AroC.

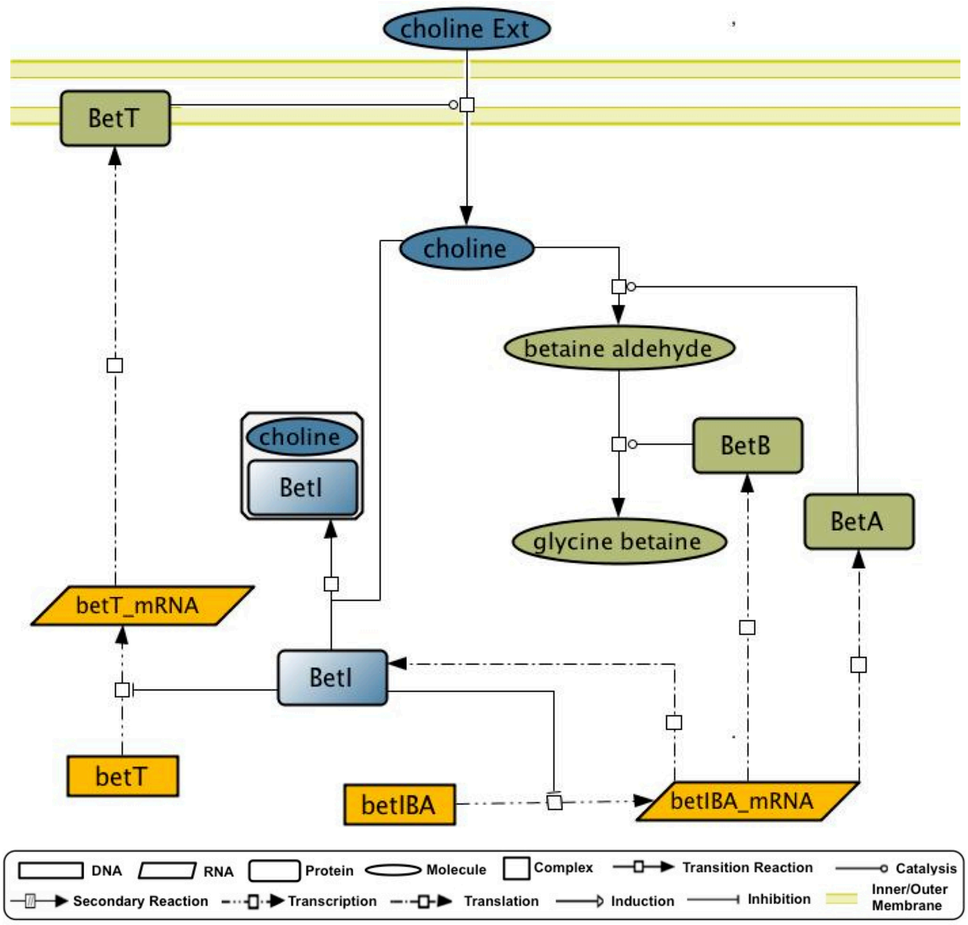


**Figure S2**. Example of GENSOR Unit. Assembled multilevel network of Transcription factor BetI. It depicts its effector, its change in conformation, regulatory effect, regulated genes and the impact they have on metabolism.

a)

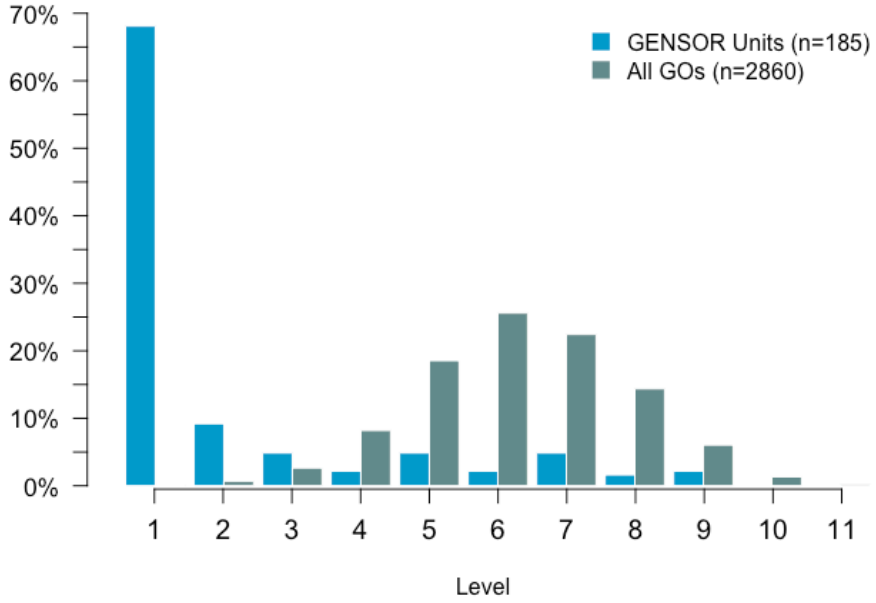
70%
60%
50%
40%
30%
20%
10%
0%
GENSOR Units (n=185)
All GOs (n=2860)
1
2
3
4
5
6
7
8
9
10
11
Level

b)

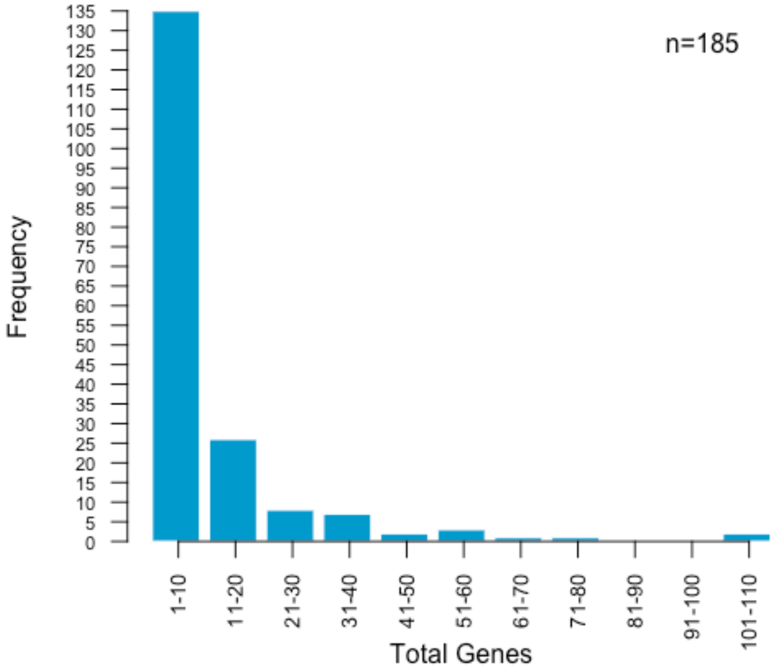
Frequency
135
130
125
120
115
110
105
100
95
90
85
80
75
70
65
60
55
50
45
40
35
30
25
20
15
10
5
0
n=185
1-10
11-20
21-30
31-40
41-50
51-60
61-70
71-80
81-90
91-100
101-110
Total Genes

c)

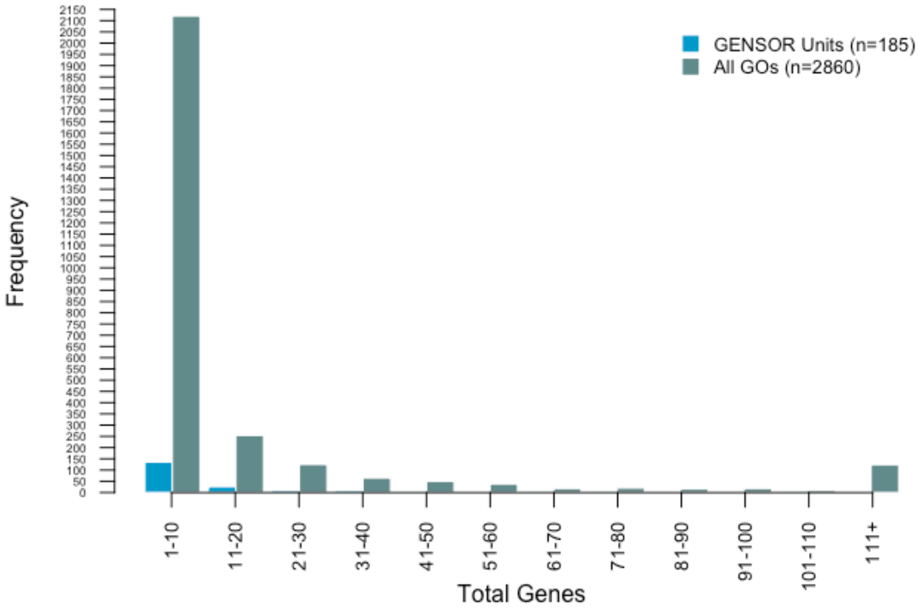
GENSOR Units (n=185)
All GOs (n=2860)
Frequency
1-10
11-20
21-30
31-40
41-50
51-60
61-70
71-80
81-90
91-100
101-110
111+
Total Genes

d)

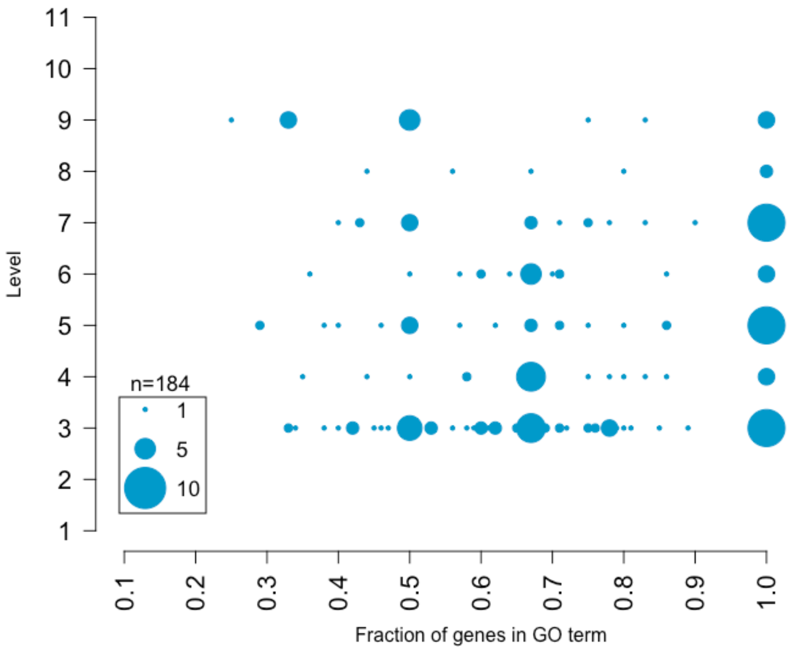
11
10
9
8
7
6
5
4
3
2
1
Level
n=184
1
5
10
0.1
0.2
0.3
0.4
0.5
0.6
0.7
0.8
0.9
1.0
Fraction of genes in GO term

e)

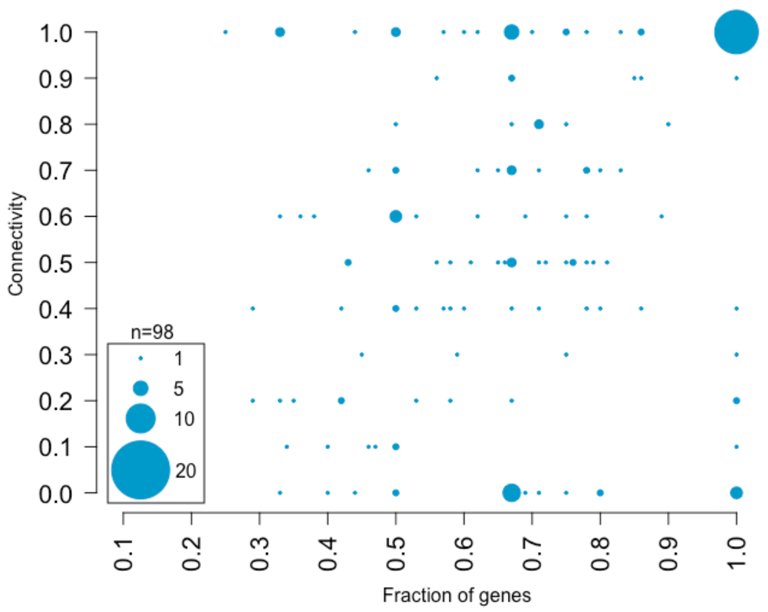


**Figure S3. (a)** Comparison of distribution of all GO terms' levels (cyan bars) with distribution of levels of representative GO terms of GENSOR Units (blue bars). **(b)** Frequencies of size of GENSOR Units. **(c)** Comparison of number of genes of GENSOR Units (blue bars) and of GO terms in level 3 and higher (cyan bars). **(d)** Comparison of the fraction of genes from a GENSOR Unit in the most representative GO term versus level of the representative GO term. Size of point reflects the number of GENSOR Units in that coordinate. **(e)** Comparison of the fraction of genes from a GENSOR Unit in the most representative GO term (excluding levels 1 and 2), versus connectivity. Size of point reflects the number of GENSOR Units. Only GENSOR Units with scores in both analyses are shown.

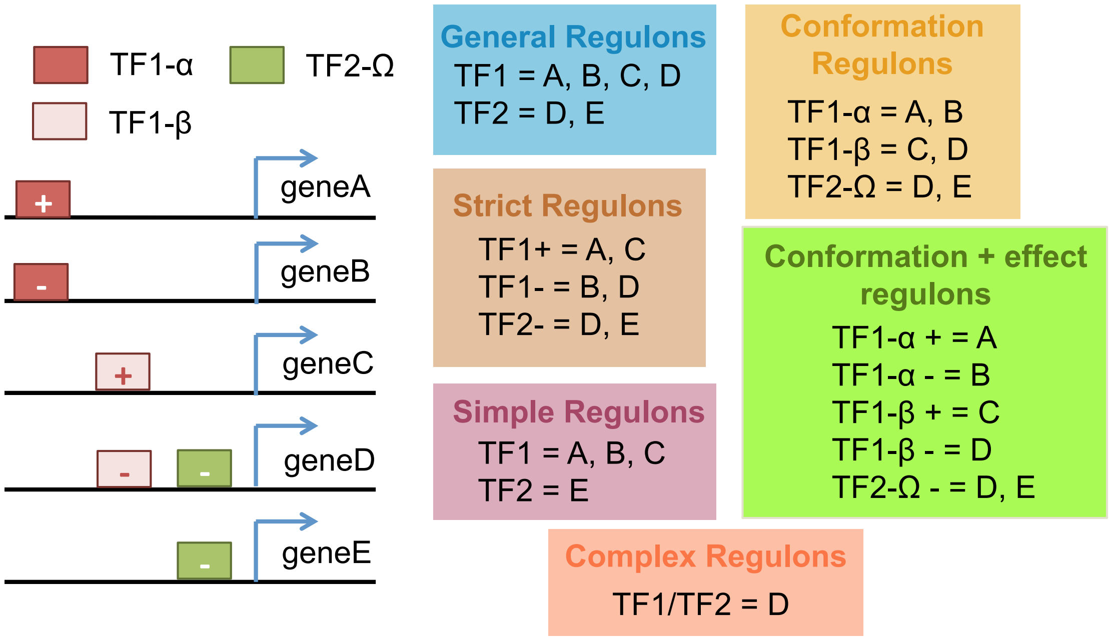


**Figure S4.** Examples of the criteria used to define regulons for each type of regulatory unit.

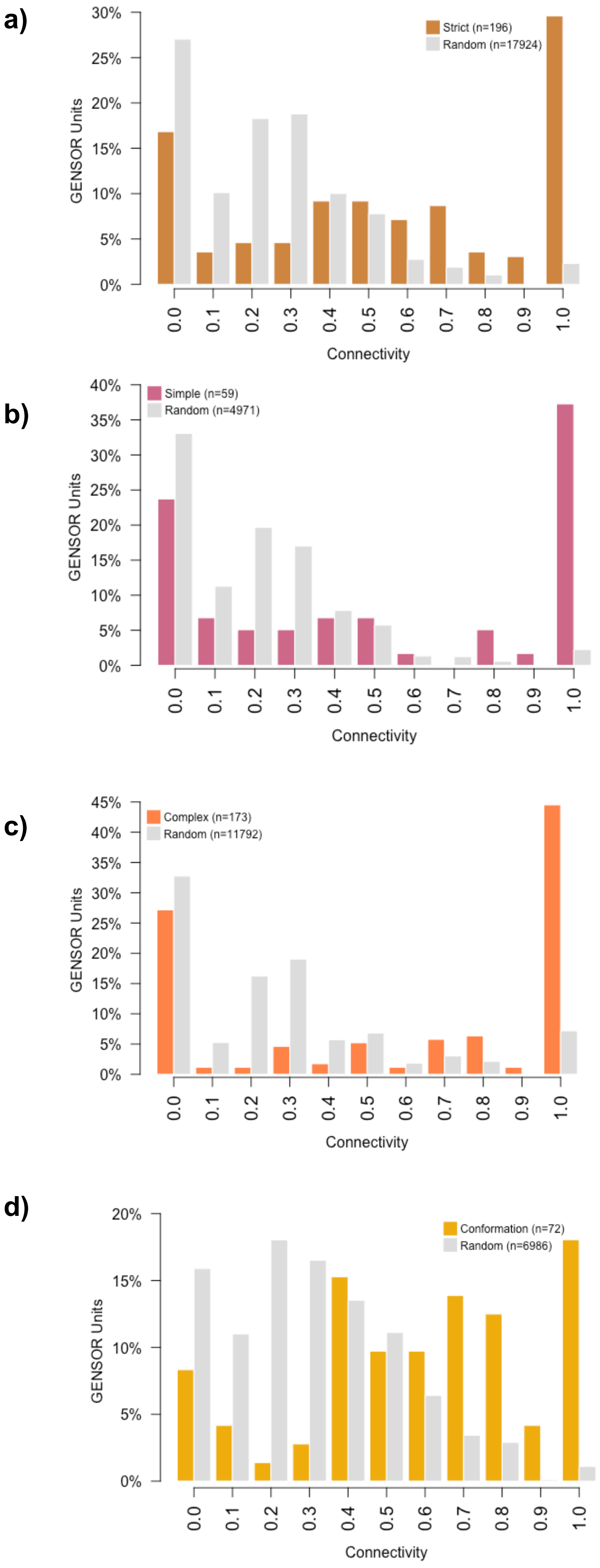
a)
Strict (n=196)
Random (n=17924)
GENSOR Units
0% 5% 10% 15% 20% 25% 30%
0.0 0.1 0.2 0.3 0.4 0.5 0.6 0.7 0.8 0.9 1.0
Connectivity
b)
Simple (n=59)
Random (n=4971)
GENSOR Units
0% 5% 10% 15% 20% 25% 30% 35% 40%
0.0 0.1 0.2 0.3 0.4 0.5 0.6 0.7 0.8 0.9 1.0
Connectivity
c)
Complex (n=173)
Random (n=11792)
GENSOR Units
0% 5% 10% 15% 20% 25% 30% 35% 40% 45%
0.0 0.1 0.2 0.3 0.4 0.5 0.6 0.7 0.8 0.9 1.0
Connectivity
d)
Conformation (n=72)
Random (n=6986)
GENSOR Units
0% 5% 10% 15% 20%
0.0 0.1 0.2 0.3 0.4 0.5 0.6 0.7 0.8 0.9 1.0
Connectivity

e)

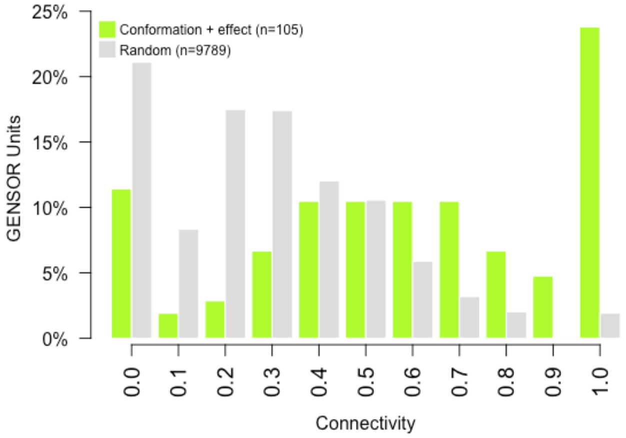


f)

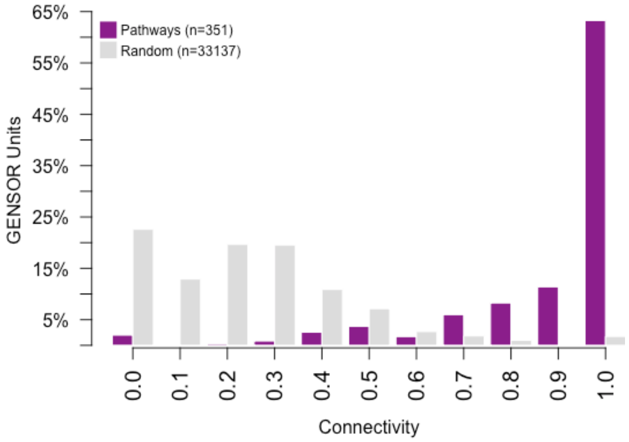


**Figure S5**. Connectivity distributions of regulatory units and pathways, compared to their random sets of regulons. All real sets were significantly different from their random counterparts (Wilcoxon-Mann-Whitney; all pvalues < 1e08). Only GENSOR Units with two or more enzymatic reactions were considered.

a)

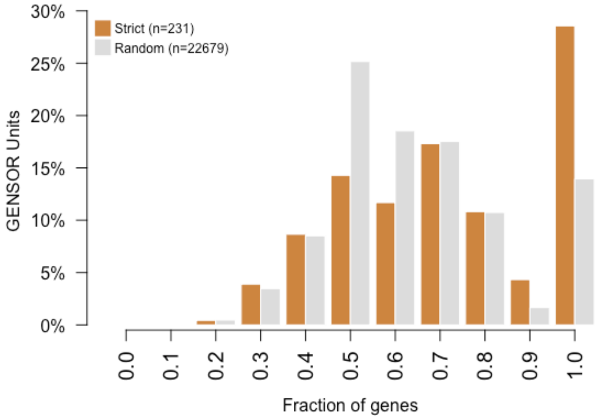

Strict (n=231)
Random (n=22679)
GENSOR Units
30%
25%
20%
15%
10%
5%
0%
0.0 0.1 0.2 0.3 0.4 0.5 0.6 0.7 0.8 0.9 1.0
Fraction of genes


b)

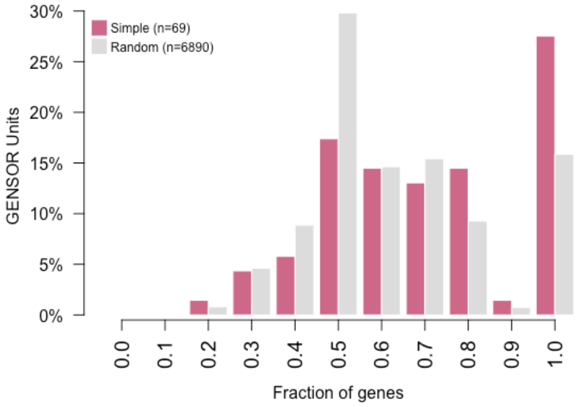

Simple (n=69)
Random (n=6890)
GENSOR Units
30%
25%
20%
15%
10%
5%
0%
0.0 0.1 0.2 0.3 0.4 0.5 0.6 0.7 0.8 0.9 1.0
Fraction of genes


c)

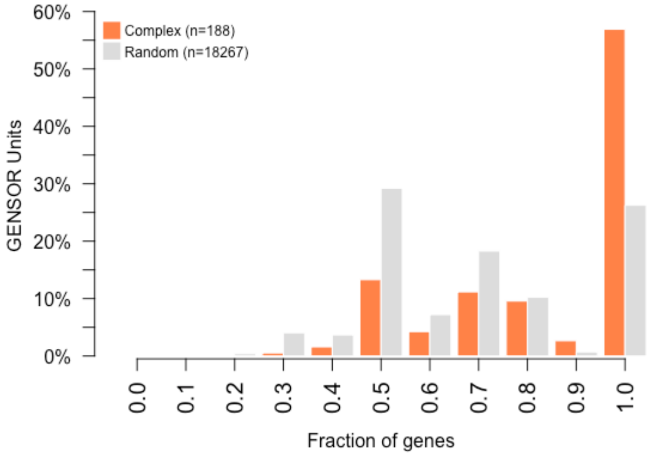

Complex (n=188)
Random (n=18267)
GENSOR Units
60%
50%
40%
30%
20%
10%
0%
0.0 0.1 0.2 0.3 0.4 0.5 0.6 0.7 0.8 0.9 1.0
Fraction of genes


d)

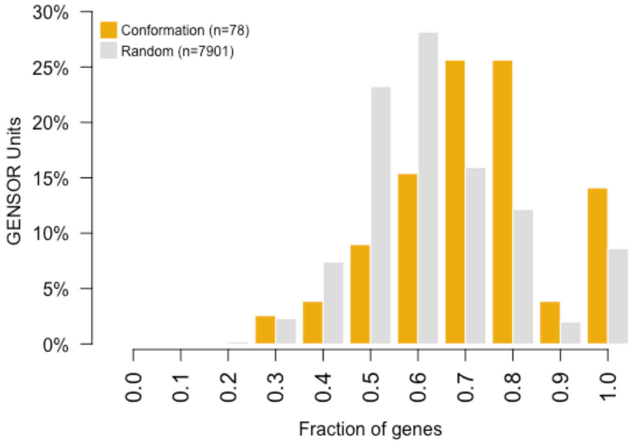

Conformation (n=78)
Random (n=7901)
GENSOR Units
30%
25%
20%
15%
10%
5%
0%
0.0 0.1 0.2 0.3 0.4 0.5 0.6 0.7 0.8 0.9 1.0
Fraction of genes

**e)**

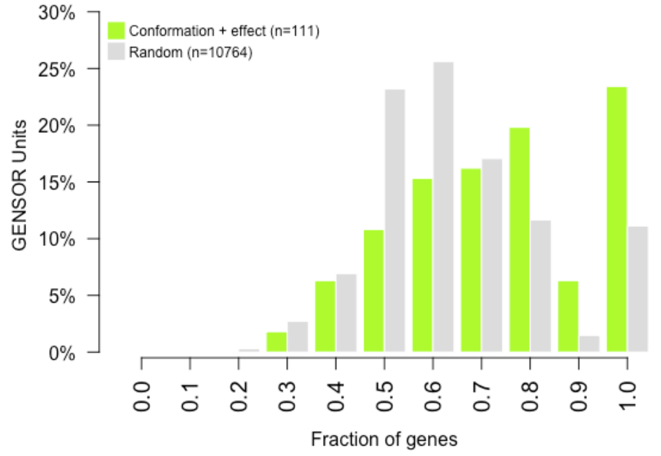


**f)**

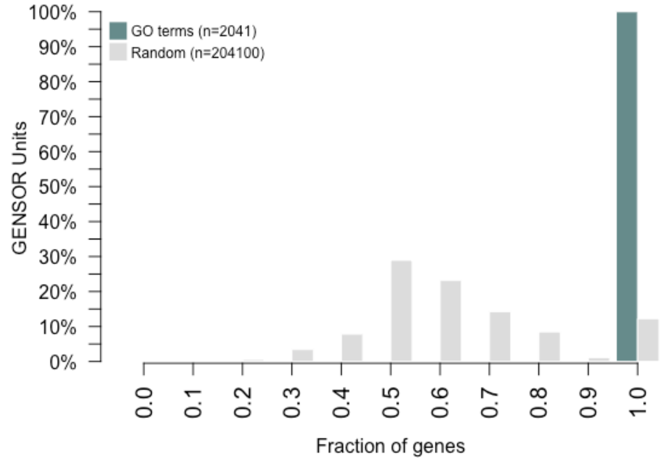


**Figure S6**. Distribution of the highest fraction of genes of each GENSOR Unit that are present in the same GO term, compared to random sets of regulons. All real sets were significantly different from their random counterparts (Wilcoxon-Mann-Whitney; all pvals < 1e03). Only descriptive GO terms (level 3 and higher) were considered. Only GENSOR Units with more than one gene annotated in the ontology were included.

| Regulatory Unit | Effect (+/-) | Regulation by other TFs | Conformation |
|---|---|---|---|
| General regulons | No | No | No |
| Strict regulons | Yes | No | No |
| Simple regulons | No | Yes | No |
| Complex regulons | No | Yes | No |
| Conformation regulons | No | No | Yes |
| Conformation + effect regulons | Yes | No | Yes |

**Table S1.** Combinations of properties considered by each regulatory unit definition.

| Regulatory units and controls | Median Connectivity / Median of random set | Median Fraction of genes in GO term / Median of random set |
|---|---|---|
| General regulons | 0.6 / 0.2 | 0.67 / 0.60 |
| Strict regulons | 0.6 / 0.2 | 0.68 / 0.60 |
| Simple regulons | 0.5 / 0.2 | 0.69 / 0.59 |
| Complex regulons | 0.8 / 0.2 | 1 / 0.67 |
| Conformation regulons | 0.6 / 0.3 | 0.70 / 0.60 |
| Conformation + effect regulons | 0.6 / 0.3 | 0.73 / 0.60 |
| Pathways | 1 / 0.2 | NA |
| GO terms | NA | 1 / 0.58 |

**Table S2**. Median scores obtained by each regulatory unit/control in the two tests for functional homogeneity.
